# High-field Functional Magnetic Resonance Imaging of Vocalization Processing in Marmosets

**DOI:** 10.1101/010561

**Authors:** Srivatsun Sadagopan, Nesibe Z. Temiz, Henning U. Voss

**Affiliations:** Laboratory of Neural Systems, The Rockefeller University, New York, NY 10065.; Department of Radiology and Citigroup Biomedical Imaging Center, Weill Cornell Medical College, New York, NY 10021.

## Abstract

Vocalizations are behaviorally critical sounds, and this behavioral importance is reflected in the ascending auditory system, where conspecific vocalizations are increasingly over-represented at higher processing stages. Recent evidence suggests that, in macaques, this increasing selectivity for vocalizations might culminate in a cortical region that is densely populated by vocalization-preferring neurons. Such a region might be a critical node in the representation of vocal communication sounds, underlying the recognition of vocalization type, caller and social context. These results raise the questions of whether cortical specializations for vocalization processing exist in other species, their cortical location, and their relationship to the auditory processing hierarchy. To explore cortical specializations for vocalizations in another species, we performed high-field fMRI of the auditory cortex of a vocal New World primate, the common marmoset (*Callithrix jacchus*). Using a sparse imaging paradigm, we discovered a caudal-rostral gradient for the processing of conspecific vocalizations in marmoset auditory cortex, with regions of the anterior temporal lobe close to the temporal pole exhibiting the highest preference for vocalizations. These results demonstrate similar cortical specializations for vocalization processing in macaques and marmosets, suggesting that cortical specializations for vocal processing might have evolved before the lineages of these species diverged.

---

The perception of vocalizations relies upon their representation in the population neural activity of neurons in the auditory pathway. In the ascending auditory system, neural activity shows a gradually increasing bias for representing conspecific vocalizations compared to other sounds. At lower processing stages of the auditory system, it is unclear if vocalizations are represented any differently than other sounds (Pollack, 2013). By the level of the inferior colliculus, however, population activity appears to over-represent conspecific vocalizations (Portfors et al., 2009; Holmstrom et al., 2010; Pollack, 2013), although only a small proportion of single neurons show selectivity for individual vocalizations (Suta et al., 2003). In early auditory cortex, single neurons start developing selectivity for features unique to conspecific vocalizations (for example, in primates, Wang and Kadia, 2001; Sadagopan and Wang, 2009). Higher in the processing hierarchy, greater selectivity for individual vocalizations is achieved by single neurons in rostral/anterior cortex (Rauschecker and Tian, 2000; Tian et al., 2001).

In macaque monkeys, this increasing representational bias for conspecific vocalizations appears to culminate in a cortical region located in the anterior auditory cortex that is densely populated with neurons that preferentially respond to conspecific vocalizations (Petkov et al., 2008, 2009; Perrodin et al., 2011). The existence of such a vocalization-selective region in macaques raises the question of whether other species exhibit similar hierarchies or cortical specializations for vocalization processing, and whether similar brain structures are involved in these specializations. In this study, we asked whether the auditory cortex of the common marmoset (*Callithrix jacchus*), a New World primate that last shared a common ancestor with the macaque lineage about 40 million years ago (Goodman et al., 1998; Steiper and Young, 2006), exhibits similar functional hierarchies for vocalization processing, and where they are localized in the brain.

Marmosets are an ideal species for investigating vocal processing because they exhibit rich vocal behaviors (Epple, 1968; Stevenson and Poole, 1976; Snowdon 2001) and possess a large, well-characterized vocal repertoire that is retained in captivity (Wang, 2000; DiMattina and Wang, 2006; Pistorio et al., 2006). The marmoset auditory system is well-studied – at the cortical level, the anatomy and connectivity of primary and higher auditory cortices are well-characterized (e.g., Aitkin et al., 1993; de la Mothe et al., 2006, 2012), basic neural response properties are known (e.g., Aitkin et al., 1986; Kajikawa et al., 2005; Philibert et al., 2005; Wang et al., 2005; Bendor and Wang, 2008), the neural basis for responses to more complex stimuli have been studied (Kadia and Wang, 2003; Sadagopan and Wang, 2009) and the neural representation of conspecific vocalizations by neurons in the primary and belt auditory cortex has been explored (Wang et al., 1995; Wang and Kadia, 2001; Nagarajan et al., 2002; Kajikawa et al., 2008). While these studies have provided a wealth of information about the initial cortical stages of vocalization processing, an open question is whether the marmoset auditory cortical pathway builds up, in a hierarchical manner, to functionally specialized cortical regions for processing conspecific vocalizations.

Scouting large swathes of cortex for such functional specializations using electrophysiology is time-consuming and constrained by the accessibility of different cortical regions for invasive experiments. As an alternative, in macaques, fMRI has been used as a powerful tool to localize such functionally specialized cortical regions, in both visual cortex – for example, in localizing face-selective cortical regions (Tsao et al., 2003, 2006), and auditory cortex – for example, in localizing vocalization-selective regions (Petkov et al., 2008). fMRI-based localizers can then be used to target electrophysiology to much smaller regions of interest (Tsao et al., 2006; Perrodin et al., 2011). In this study, we used high-field fMRI to investigate the activation of marmoset auditory cortical areas by complex auditory stimuli. We demonstrate the existence of a caudal-rostral gradient for preferential vocalization processing in marmosets, with rostral regions close to the temporal pole exhibiting the maximal preference for conspecific vocalizations. Our results demonstrate that similar structure-function relationships might operate in marmosets and macaques for vocalization coding, and provide a basis for detailed studies of marmoset temporal pole regions using electrophysiological and high-resolution imaging methods to probe the neural basis of the processing of vocal communication sounds.

## Results

The goals of this study were to determine if functional specializations for vocalization processing exist in the auditory cortex of common marmosets, and to determine the relationship of these regions to the auditory processing hierarchy. To this end, we used fMRI to measure BOLD activity in the auditory cortex of anesthetized marmosets using both vocalizations and simpler (tone) stimuli. Our main finding is the existence of a caudal-rostral gradient for vocalization processing, with anterior temporal lobe regions located close to the temporal pole (TP) and rostro-lateral to tone-responsive cortex having the greatest selectivity for vocalizations. BOLD data were acquired in a 7.0 Tesla small animal scanner, in which marmosets were placed by non-invasively securing them to a custom anatomical positioner (Figure 1). We emphasize the non-invasive nature of the restraint because this helped reduce the required level of isoflurane anesthesia, which proved to be a critical determinant of auditory cortex responsivity.

**Figure 1:**
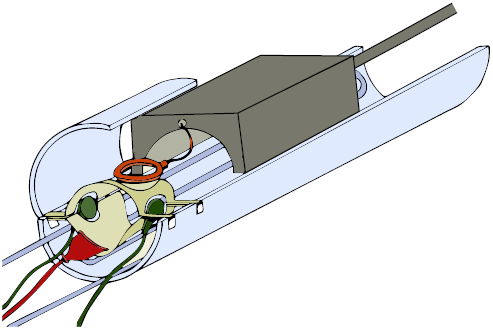
Experimental setup for marmoset auditory fMRI. The marmoset was placed, under anesthesia, in an anatomical positioner with built-in warm water heating support (blue) using non-invasive restraints and acoustic isolation foam. A custom 3D printed, foam-lined helmet (beige) was used to restrain the marmoset’s head with ring coil (orange). Isoflurane anesthesia was delivered through a face mask (red), and acoustic stimuli were presented through MR-compatible earphones (green). The coil preamplifier box (gray) acted as a restraint for the subject’s body.

### Sparse imaging paradigm

To enable the delivery of acoustic stimuli with minimal interference from scanner noise, we used a sparse slice acquisition paradigm (Figure 2; similar to Petkov et al., 2009). In this paradigm, we first restricted the imaged area to six 1.2 mm-thick slices positioned obliquely and parallel to the lateral sulcus (LS), covering the expected location of auditory responsive areas (Figure 3A). Because of the small number of slices acquired, data acquisition time and the concomitant gradient-switching noise were brief (<0.25 s). We acquired one complete volume every 2.25 s (red lines and regions in Fig. 2), allowing us a ∼2 s period of low ambient noise during which acoustic stimuli could be presented. In this way, close to 90% of acoustic stimulation was presented free from masking by scanner noise. We adopted a block design for the experiments with alternating ON and OFF blocks, each 22.5 s long, with the ON blocks consisting of either 1) vocalization or 2) tonal stimuli. In the vocalization experiment, we used three different ON blocks (Fig. 2A) consisting of conspecific vocalizations (V; Fig. 2B, top), phase-scrambled vocalizations (N, for ‘noisy’) and heterospecific vocalizations (H). In the tone experiment, we used two ON blocks consisting of high-frequency (Fig. 2B, bottom) and low-frequency tone pips (see Materials and Methods for details).

**Figure 2:**
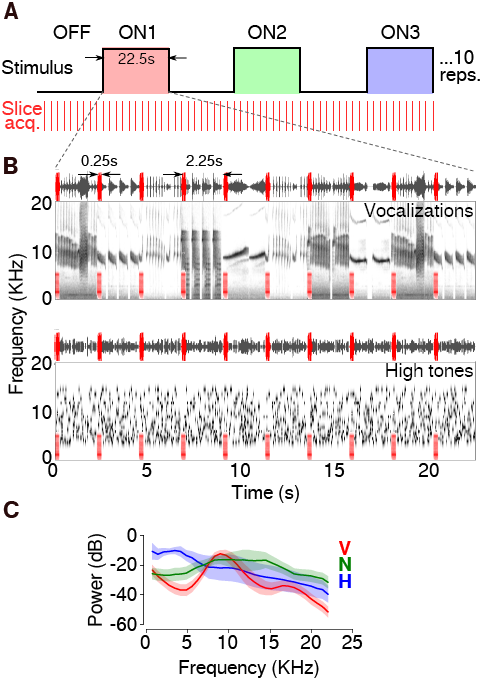
Sparse acquisition of auditory BOLD responses. (A) Stimuli in each run were organized into alternating OFF and one of three ON blocks, each 22.5s in length, repeated 5 – 10 times. Data acquisition was triggered every 2.25s (red lines), with the scanner requring <= 0.25s to acquire data from 6 slices. Red, blue and green shading correspond to the three stimulus types used in the vocalization experiment – conspecific vocalizations (V), phase-scrambled vocalizations (N) and heterospecific vocalizations (H). Tone experiments had two ON blocks – low frequency and high frequency tones. **(B)** Waveform and spectrogram of vocalization (top) or high frequency tone (bottom) stimuli contained in one ON block. Overlaid in red are waveforms and spectrograms of scanner noise at times of slice acquisition. **(C)** Average power spectra (shading corresponds to 1 s.e.m.) for the three stimulus categories used in the vocalization experiment (red – V; green – N; blue – H).

### Vocalization experiment

Thirty experimental runs, each run consisting of ten repetitions of three stimulus blocks, were acquired for the vocalization experiment. In 18 of these 30 runs, the isoflurane level could be kept below 1%, without notable movement artifacts (see Materials and Methods). We could not elicit significant BOLD responses in the remaining runs, in which isoflurane level was above 1%. Analyzing the average activation of auditory cortex, we discovered significant bilateral auditory cortex activation in 6 runs and unilateral activation in a further 6 runs (false discovery rate with cluster size thresholding; FDR-corrected q-value = 0.05; cluster threshold, 8 contiguous-face voxels). We used only the 6 runs (from 4 subjects) in which we obtained bilateral activation for further analysis in the vocalization experiment.

In Figures 3A and 3B, we show how the imaged slices and the regions of auditory-evoked activation in the vocalization experiment were positioned relative to the whole brain. In this example (Subject C), we observed significant bilateral activation of regions close to the LS, extending ∼6 mm deep from the lateral surface of the brain at its maximal extent. We also observed activation of midbrain auditory regions in this subject, but this activation was not reliably repeatable across subjects. We plotted the average activation in response to all three stimulus types overlaid on a high-resolution anatomical image to better localize these regions of activation. In Figure 3C, the heatmap corresponds to the magnitude of BOLD signal change from baseline, the transparency corresponds to the absolute value of the t-statistic, and the black contour corresponds to t ≥ 5 (Allen et al., 2012). Activation of lateral temporal lobe close to LS, as well as midbrain regions, was evident. We then extracted the time-course of the BOLD signal averaged across all voxels that passed the t ≥ 5 threshold (Fig. 3D) to visualize BOLD activation throughout the duration of the experiment. We further averaged the time-course over all repetitions to obtain the average activation for each of the three stimulus types, across all significant voxels in the cortex (Fig. 3E). In this subject (Subject C), the average magnitudes of the BOLD responses (rel. to baseline) were 0.76% (conspecific vocalizations), 0.93% (phase-scrambled vocalizations) and 1.15% (heterospecific vocalizations) – a grand-average BOLD response (across all blocks) of 0.95% signal change relative to baseline. Over all 6 runs (4 subjects) used for analysis, the grand average BOLD response was 0.49% signal increase over baseline. The peak magnitude of auditory-evoked BOLD activation in our experiments was comparable to that elicited in the somatosensory cortex of anesthetized marmosets by electrical stimulation of the forearm (Liu et al., 2013).

**Figure 3:**
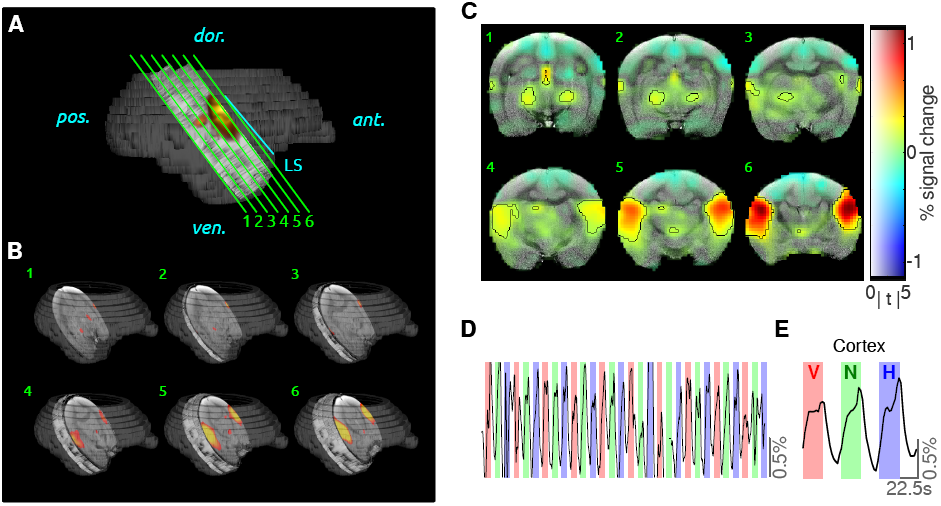
BOLD activity in the auditory cortex. (A) Six slices (green lines) of 1.2mm thickness were positioned parallel to the lateral sulcus (LS; blue line), with the last slice abutting the LS. Dark grey structure is a surface reconstruction of Subject C’s brain from an anatomical scan, light gray region corresponds to the indexed location of functional slices, and heatmap corresponds to regions of significant BOLD activation (t ≥ 5). **(B)** Three dimensional view showing slice positioning relative to the whole brain, and regions of significant BOLD activation (heatmap). **(C)** Average predictor value (beta) for the three vocalization stimuli mapped on to an anatomical MRI (left side of figure is left hemisphere). Here the heat map corresponds to the percent change in BOLD activation, transparency corresponds to the t-statistic, and black contours encompass regions with a t-statistic greater than five (corresponds to FDR-corrected q value < 10-5). **(D)** Average time-course from all voxels with t>5, across 10 repetitions of the three vocalization stimuli – conspecific vocalizations (V; red shading), phase-scrambled vocalizations (N; green shading) and heterospecific vocalizations (H; blue shading). **(E)** Time-course data in (D) averaged over all 10 repetitions for significant cortical voxels (t>5).

We then asked whether distinct regions of the brain were differentially activated by the three types of vocalization stimuli. To determine this at the level of single subjects, we first plotted contours corresponding to regions of significant brain activation (FDR corrected, q = 0.05) for each stimulus type. In Figure 4A, we plot these contours for the same subject (Subject C) as in Figure 3. We then summarized these activation patterns in an anatomically-referenced matrix as follows. First, we defined a 24 × 6 matrix per hemisphere, starting at the temporal pole and extending along the LS for 15 mm (0.625 mm bins), and starting at the LS and extending 7.2 mm lateral to the LS (1.2 mm bins; magenta boxes in Fig. 4A show one column of the complete matrix). Each element of this 24 × 6 matrix was the average of the beta values derived from a 4 voxel deep area (each magenta box). Figure 4B plots this matrix derived from Subject C. While we observed a greater number of rostral matrix elements active for conspecific vocalizations compared to phase-scrambled vocalizations, contrasting these two stimulus types at the single-subject level did not yield statistically significant differences. The number of active matrix elements was similar for conspecific and heterospecific vocalizations. Thus, the singe-subject data neither confirm nor exclude regional vocalization preferences.

**Figure 4:**
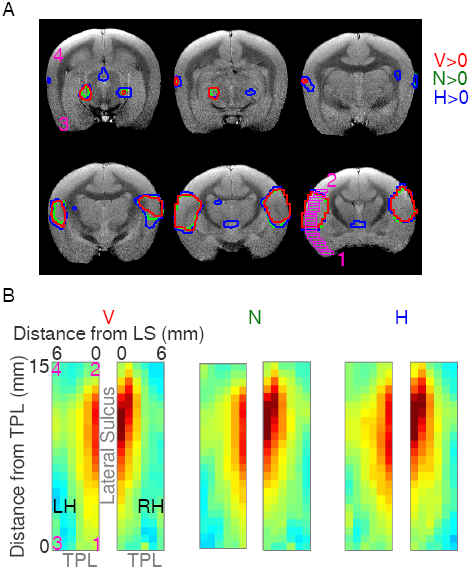
Auditory BOLD activation by different stimulus categories in a single subject. (A) Data from Subject C separated by stimulus type. Colored contours on the anatomical images correspond to regions of cortex significantly activated (t>5) by: red – conspecific vocalizations (V), green – phase-scrambled vocalizations (N) and blue – heterospecific vocalizations (H). **(B)** Auditory cortex activation in each subject can be summarized in the form of a matrix, extending 15mm caudally from the temporal pole (TPL) and 7.2mm laterally from the lateral sulcus (LS). Numerals in magenta indicate the anatomical locations to which the corners of the matrix correspond to, and each element in the matrix contains the beta value corresponding to each stimulus type averaged across 4 voxels (magenta boxes in A). Matrix on the left corresponds to the left hemisphere (LH). Color map corresponds to the normalized beta value.

To obtain more statistical power to assess if functionally specialized cortical regions existed, we performed group analysis across all scanned subjects. Because the brain sizes of the scanned subjects were similar, and because slice position was consistently determined by anatomy, we could then use the activation matrices across subjects to determine the average activation pattern, referenced to anatomical landmarks, for each stimulus type. In Figure 5A, the group-averaged activation maps are plotted, and matrix elements with black outlines correspond to those regions that were significantly higher than baseline (FDR corrected, q ≤ 0.05). We noticed that, compared to phase-scrambled vocalizations, a greater number of rostral and rostro-lateral elements of the matrix were active for conspecific and heterospecific vocalizations. To statistically evaluate preferential cortical processing of vocalizations, we then performed second-level GLM analyses contrasting conspecific vocalizations against the other stimuli. These results will be discussed later in this section.

**Figure 5:**
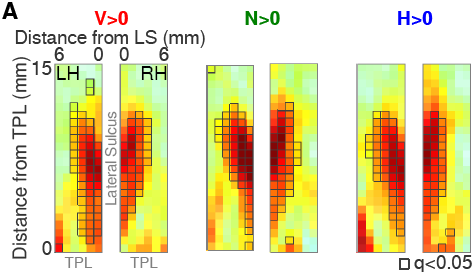
Average BOLD responses to vocalization stimuli across 4 subjects. (A) Activation summary matrices were averaged across 6 imaging runs from 4 subjects. Colormap corresponds to normalized average beta values. Matrix elements with gray outlines are those that display significant activation (FDR-corrected, q<0.05) by that stimulus type compared to baseline. A greater number of rostral voxels located close to the temporal pole appear to be activated by conspecific vocalizations.

### Tone experiment

To functionally localize areas of activation by complex stimuli relative to tone responses, we performed single-subject and group analyses as above for 5 runs of the tone experiment (out of 20 runs) from three subjects in which we elicited significant bilateral tone activation. In each of the three subjects, we observed alternating regions of cortex that were responsive to low- and high-frequency tones, as expected for tonotopically organized cortex (Figure 6A). Because the precise mapping varies from subject to subject, direct averaging across subjects (as performed in Figure 5) would systematically underestimate tonotopy at the group level. Therefore, we performed a second-level general linear model (GLM) analysis on the matrices obtained from individual subjects, with predictors for stimulus type and subject. We treated the two hemispheres of each subject independently to increase statistical power for these analyses. We then determined which elements of the response matrix showed significant second-level beta values for either stimulus type (Figure 6B). By doing so, we were able to define the cortical regions which were, on average, tone-responsive (matrix elements with magenta outlines). In the section below, we use this information in conjunction with the vocalization experiment to localize regions of cortex that show differential activation by vocalizations relative to tone-responsive cortex.

**Figure 6:**
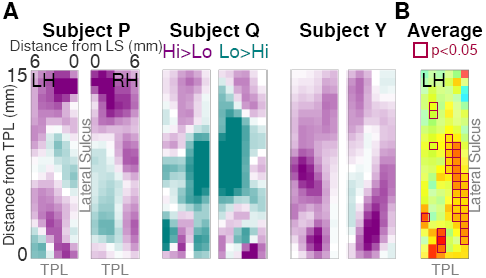
Activation of auditory cortical regions by tone stimuli. (A) In each imaged subject, we observed an alternating spatial pattern of cortical responses to high- and low-frequency tones. Magenta regions correspond to cortical regions that were better activated by high-frequency stimuli, and cyan regions were better activated by low frequency stimuli. The central cyan region of activation corresponds to the low-frequency regions of A1 and R. **(B)** Map of beta values responding significantly to tones of any frequency. Matrix elements with magenta borders are significantly activated by tones compared to baseline (p<0.05; not corrected for multiple comparisons).

### Differential activation of rostro-lateral auditory cortex by conspecific vocalizations

To determine if there exists a preference for conspecific vocalizations in some cortical regions, we performed second-level GLM analysis as above for the vocalization dataset in Figure 5, also treating hemispheres independently to increase statistical power. When we contrasted conspecific vocalizations against phase-scrambled vocalizations (V>N, Fig. 7A, left) or against heterospecific vocalizations (V>H, Fig. 7A, right), we observed that rostral and rostro-lateral regions of cortex, close to the temporal pole, exhibited a preference for conspecific vocalizations, with the most rostro-lateral region of imaged cortex exhibiting the highest preference (0.2% signal change from baseline; p=0.03, not corrected for multiple comparisons; green arrows in Fig. 7). Overall, the magnitudes of the differential activation in the rostral regions were about 0.08% signal change from baseline. Because the magnitude of the average BOLD response across stimulus categories was about 0.5%, the observed 0.2% maximal signal change and 0.08% average signal change correspond to a 40% and 16% increase in the response magnitude for conspecific vocalizations relative to phase-scrambled vocalizations. The differences in magnitude were about half as much for conspecific versus heterospecific vocalizations and were not statistically significant in individual matrix elements. We emphasize here the spectral similarity of conspecific (V) and phase-scrambled (N) vocalization stimuli – because the phase-scrambled stimuli were generated from the same set of conspecific vocalization tokens used in the experiments, they had highly overlapping average spectra, and differed mostly in their higher-order spectrotemporal structures. Thus, the observed differences in BOLD responses between V and N is a result of an underlying cortical process that is sensitive to the spectrotemporal features of conspecific vocalizations. For illustration, we have plotted in Figure 7B the stimulus category that elicited the best response in each matrix element. From this plot, one can observe the transition of stimulus preference from noisy stimuli in caudal auditory cortex to spectrotemporally complex stimuli in rostral auditory cortex, with conspecific vocalizations being the optimal category near temporal pole.

**Figure 7:**
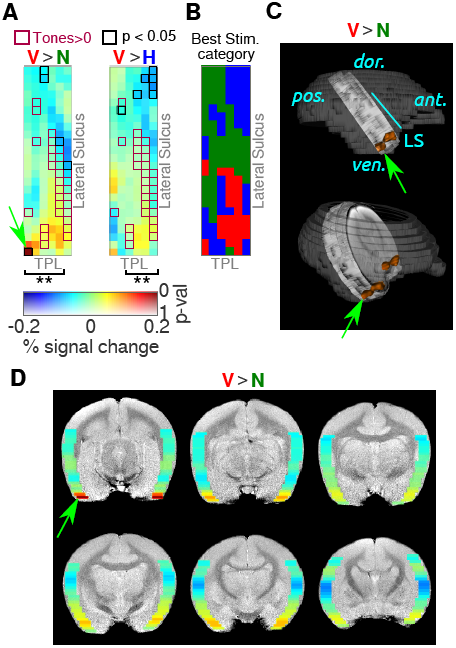
A caudal-rostral gradient for vocalization selectivity in auditory cortex. (A) Differential response map of cortical activation derived from second-level GLM analysis. Warmer colors of the heat map correspond to regions that were better activated by conspecific vocalizations, compared to phase-scrambled (V>N; left) or heterospecific vocalizations (V>H; right). Transparency of the heatmap corresponds to uncorrected p-value. Matrix elements outlined in magenta are tone-responsive regions (from Fig. 6). Those outlined in black are statistically significant (p<0.05; not corrected for multiple comparisons). Asterisks denote columns which exhibited a significant caudal-rostral gradient in vocalization selectivity (**: q ≤ 0.01, where q is the Bonferroni corrected p-value of a linear regression between the differential response magnitude and caudal-rostral location). Only left hemispheres are shown because we treated the hemispheres independently for group analysis. **(B)** Stimulus category eliciting the best response in each matrix element. Colors denote conspecific vocalizations (V; red), phase-scrambled vocalizations (N; green) and heterospecific vocalizations (H; blue). **(C and D)** Differential response map projected back into anatomical coordinates to illustrate the caudal-rostral gradient and location of the most vocalization-selective part of the gradient. Green arrow and orange regions correspond to regions most selective for conspecific vocalizations.

We tested if there exists a gradient in the preference for conspecific vocalizations along the LS by fitting a line to the differential effect size along the caudal-rostral direction (along columns of the matrix in Fig. 7A). We found that preference for conspecific vocalizations increased in the caudal to rostral direction along the lateral sulcus. Statistical significance was evaluated from the p-value of a linear regression on each column of the group average matrix (corresponding to slices along LS), Bonferroni-corrected for multiple comparisons (asterisks in Fig. 7A correspond to corrected p ≤ 0.01). The R^2^ and Bonferroni-corrected p-values for the linear regression on significant V>N columns in Fig. 7A (left; going from left to right) were R^2^ = 0.39 (p = 0.01), 0.43 (0.003), 0.88 (5.2 × 10^-11^), 0.67 (6.6 × 10^-6^) and 0.63 (2.5 × 10^-5^). Similarly, R^2^ and Bonferroni-corrected p-values of the significant V>H columns (left to right) were R^2^ = 0.51 (p = 0.0006), 0.83 (5 × 10^-9^), 0.91 (2.2 × 10^-12^) and 0.57 (1.1 × 10^-4^). Functionally, the cortical regions that exhibited a greater preference for conspecific vocalizations were not significantly tone-responsive (magenta boxes in Fig. 7A), and located rostral and rostro-lateral to tone-responsive cortex. Figure 7C is a remapping of the V>N matrix in Fig. 7A onto a high-resolution anatomical scan – the caudal-rostral gradient in vocalization selectivity is readily apparent. In Fig. 7D, we illustrate the anatomical location of the anterior end of this gradient, where we would expect the most vocalization-selective responses.

Compared to conspecific or heterospecific vocalizations, i.e., stimuli with rich spectrotemporal structure, the cortex caudal to tone-responsive regions was better driven by phase-scrambled (noisy) stimuli (blue regions in Fig. 7A). Maximally, caudal regions exhibited about a 20% increased response to noisy sounds compared to conspecific vocalizations. This preference for broadband stimuli is consistent with broader tuning bandwidths that have been observed in caudal auditory cortex in macaques (Recanzone et al., 2000) and marmosets (Kajikawa et al., 2005; Zhou and Wang, 2014). Averaged across all slices (average of all columns of the matrix in Fig. 7A), the caudal-rostral gradient changed from an 8% preference for noisy sounds caudally to a 13% preference for conspecific vocalizations rostrally (R^2^ = 0.77, corrected p = 1.4×10^-7^). At the individual slice level, the extrema of observed values were about a 20% increased response to noisy sounds caudally (close to LS) and 40% increased response for conspecific vocalizations rostro-laterally (close to temporal pole and furthest from LS).

As a control to confirm the statistical validity of the caudal-rostral selectivity gradients for V>N and V>H, we also performed a permutation test as follows. The differential effect size matrices in Fig. 7A were randomly rearranged 100,000 times, and the R^2^ and p-value of linear fits along the columns calculated. From these randomizations, we computed the likelihood of observing a caudal-rostral gradient at the minimal R^2^ and p values observed in the data, so that the resultant probabilities provided an upper bound for how likely it was to find a gradient purely by chance. We found that for the conspecific versus phase-scrambled case (V>N), the probability of obtaining an R^2^ ≥ 0.39 (the minimum of significant R^2^ values in the data) in any single column with a significance value of p ≤ 0.0018 (the minimum uncorrected significance value in the data) was P(V>N) = 4.5 × 10^-4^. Similarly, for conspecific versus heterospecific vocalizations (V>H), the probability of obtaining a single-column gradient with R^2^ ≥ 0.51 and significance value p ≤ 1.2 × 10^-4^ was P(V>H) = 3.7 × 10^-5^. Simultaneous gradients along multiple columns did not occur even once over the 100,000 randomizations for both the V>N and V>H comparisons, whereas in the data we observed five and four simultaneous gradients respectively. The above test assumed independent matrix elements, and in each permutation, the actual numerical values of the matrix elements were preserved. A more stringent control would be to preserve spatial correlations between the matrix elements; this can be effected by computing the 2D spectrum of the matrices, retaining the power spectrum, and scrambling only the phases. This computation, however, comes with the trade-off of altering the numerical values of individual matrix elements in each permutation. When we repeated the permutation test with phase-scrambled matrices as outlined above, the probability of observing single-column gradients at the minimum observed levels in the data were: P(V>N) = 0.058 and P(V>H) = 0.023. The probability of observing 5 simultaneous gradients in the V>N case was 4.9 × 10^-4^, and that of observing 4 simultaneous gradients in the V>H case was 5.0 × 10^-5^. Therefore, we conclude that the observed caudal-rostral gradient in conspecific vocalization selectivity was a statistically highly significant and non-random arrangement.

## Discussion

These data demonstrate a gradient for the preferential processing of conspecific vocalizations in a caudal-rostral direction along the LS, with rostro-lateral cortical regions close to the temporal pole exhibiting the most preference for conspecific vocalizations. Regions that exhibited the most preference for vocalizations appeared to lie outside of tone-responsive cortex. Our study points to a homologous structure-function relationship in the processing of conspecific vocalizations in the brain between a New World primate (the common marmoset) and an Old World primate (the macaque), suggesting that cortical specialization for vocalization processing may have evolved before the lineages of these two species diverged from a common ancestor ∼40 million years ago (Goodman et al., 1998; Steiper and Young, 2006). Finally, our study proposes a target for electrophysiological studies of marmoset vocal processing.

### Homologous structure-function relationships of cortical processing in primates

In macaque monkeys (in both awake and anesthetized animals), fMRI experiments have suggested that a ∼50mm^3^ cortical region selective to conspecific vocalizations and individual identity, situated in the anterior temporal lobe close to the temporal pole, lies at the apex of the auditory processing hierarchy (Petkov et al., 2006, 2008, and 2009). The differential magnitude of vocalization responses observed in these studies was ∼0.6% BOLD signal change. In the present study, we did not find evidence for well-defined cortical regions that preferentially processed conspecific vocalizations. We do find, however, a similar anatomical pattern for encoding conspecific vocalizations in marmosets – anterior temporal lobe regions lie at the upper end of a vocalization-selectivity gradient, with a maximal differential magnitude of about 0.2% BOLD signal change. Thus the propensity of rostral temporal lobe regions in marmosets to preferentially process vocalizations is consistent with the organization of macaque auditory cortex.

If a marmoset vocalization region did exist, it is worth considering what the expectation for the size of such a region should be. For a rough estimate, let us make the simplifying assumption that the volume of functionally specialized areas scales linearly with the total volume of gray matter in the cerebral cortex. Marmosets have lissencephalic brains with a neocortical volume of ∼4400 mm^3^, whereas the brains of macaques are highly gyrencephalic with a neocortical volume of ∼63500 mm^3^ – 14.5 times bigger than marmosets (Dunbar, 1992; Chaplin et al., 2013). A 50 mm3 volume (size of the Petkov voice region) of functionally specialized cortex in macaques would therefore scale to a ∼3.5 mm^3^ volume in marmosets – a sphere of radius 0.94 mm, or, at our imaging resolution of 0.625 × 0.625 × 1.2 mm, about 7 voxels. Accounting for nonlinearities in the expansion of different cortical areas, it appears that auditory cortical regions and temporal pole has expanded between 8x – 16x in macaques compared to marmosets (Chaplin et al., 2013), suggesting estimates of between ∼3 – 6 mm^3^ (6 – 13 voxels) for a putative vocalization area in marmosets. This small expected size might have therefore rendered such a specialized cortical region difficult to detect. For example, a recent study comparing fMRI maps to maps derived from high resolution neurophysiology data suggested that fMRI was most useful for detecting large (> 2.5 mm) domains of selective cortex (Issa et al., 2013). What we observe as a gradient in vocalization selectivity might therefore be a spatially-smoothed reflection of an underlying cortical specialization. Alternatively, compared to macaques, vocalization-preferring neurons in marmosets may be more diffusely distributed in the anterior auditory cortical regions.

### The rostral auditory cortex and the auditory processing hierarchy

Convergent lines of evidence point to the rostral regions of auditory cortex being more selective for conspecific vocalizations in primates – for example, in macaques, neurons in the AL belt area exhibit high vocalization selectivity (Tian et al., 2001), regions in the anterior auditory cortex are populated by more vocalization-selective neurons (Perrodin et al., 2011) and discrimination of certain categories of vocalizations is enhanced in rostral auditory cortex (Fukushima et al., 2014). Taken together, these data suggest that rostral regions of the temporal lobe in macaques display response characteristics that are ideally suited for processing sounds with complex spectrotemporal patterns such as vocalizations (Romanski and Averbeck, 2009). Thus, the caudal-rostral gradient that we observe for vocalization processing in marmosets is also consistent with a range of electrophysiological studies in macaques, and suggest that anterior temporal lobe regions in marmosets are at the apex of the sensory auditory processing hierarchy as well.

In broader terms, increasing selectivity for vocalizations has been observed in other along an anterior/ventral direction. In evolutionarily earlier species such as Guinea pigs, there is some experimental evidence that rostral and ventral secondary cortical areas respond to spectrotemporally complex sounds such as vocalizations, whereas caudal and dorsal regions do not respond to vocalizations (Grimsley et al., 2013). In dogs, Andics et al. (2014) have reported that anterior and ventral auditory cortical areas are maximally responsive to dog vocalizations. In more recently evolved species such as humans, imaging experiments have demonstrated communication sound selective regions in the anterior temporal lobe (Scott et al., 2000; Kriegstein and Giraud, 2004) and macaques (Petkov et al., 2008), but also in roughly corresponding brain regions of evolutionarily more distant species such as dogs (Andics et al., 2014). We note, however, that the anterior pathway may not be the only vocalization-selective pathway in higher primates. For example, in macaques, a communication sound preferring region of the insula has been reported (Remedios et al., 2009). In chimpanzees, a PET imaging study suggested that preference for conspecific vocalizations is localized to the posterior temporal lobe (Taglialatela et al., 2009). In humans, conspecific vocal sounds activate central and posterior regions of the superior temporal sulcus in addition to anterior regions (Belin et al., 2000). In our data, however, we did not find any caudal vocalization-selective regions or reversals of the vocalization-selective gradient at a caudal location. Our study supports the importance of anterior auditory cortex in the perception of vocal communication sounds.

### Auditory fMRI under anesthesia

One limitation of the present study is that we conducted our experiments under anesthesia. The primary reason for performing our experiments under anesthesia was to keep motion artifacts as minimal as possible using non-invasive techniques – because of the partial slice prescription necessitated by the sparse scanning paradigm, any motion artifact that resulted in out-of-slice motion was irrecoverable. A second reason was to develop non-invasive techniques of obtaining fMRI data, which would enable the collection of comparative data from a variety of evolutionarily interesting species, providing crucial insight into brain evolution. But anesthesia is a critical determinant of the response characteristics of auditory cortical neurons, with effects on response magnitude (Wang et al., 2005), latency and reliability (Ter-Mikaelian et al., 2007), transience (Wang et al., 2005) and tuning properties (Gaese and Ostwald, 2001). Keeping these effects in mind, we took great care to keep the level of anesthesia as low as possible in these non-invasive experiments; by introducing nitrous oxide into the gas mix, we were able to image at isoflurane levels of 0.25 – 0.75%. Even the low levels of anesthesia we used might have altered some underlying neural response properties; perhaps inducing the biphasic hemodynamic response function that we observed (Materials and Methods). But at the same time, anesthesia has not seriously hindered the localization of cortical specializations in previous fMRI studies in macaques: anesthetized functional localization (using remifentanil) of both higher auditory (Petkov et al., 2008) and higher visual (Ku et al., 2011) cortex are similar to that observed in awake animals. Robust, bilateral auditory BOLD responses can be obtained under anesthesia (intravenous ketamine and isoflurane) in cat auditory cortex (Hall et al., 2014). In marmoset somatosensory cortex, the primary effect of propofol anesthesia appears to be about a 50% suppression of BOLD response magnitude (Liu et al., 2013). In rats, differential cortical responses to auditory stimuli, including human voices, have been observed under deep isoflurane anesthesia (Rojas et al., 2008). Thus, while it is possible that we may have missed strong responses in higher auditory cortical regions in our experiments, and consequently missed extant small, specialized cortical regions, we expect the observed caudal-rostral gradient in vocalization selectivity to generalize well to awake animals.

There are many improvements in the experimental preparation that could increase the power of fMRI imaging of auditory cortex in marmosets. First among these is imaging awake marmosets, which can be accomplished with the use of custom 3D-printed helmets (for example, Liu et al., 2013) or MR-compatible headposts for head fixation. However, as mentioned earlier, for sparse-scanning paradigms, great care should be taken to minimize motion artifacts. A second technical advance is the use of multichannel RF coil arrays (Papoti et al., 2013) that could result in increased whole-brain coverage with high signal-to-noise ratio. It is unclear if having the animals perform an active task during imaging would result in a better signal-to-noise ratio – for example, in macaques, behavior does not confer an advantage for spatial discrimination (Scott et al., 2007), and in rodents, neural responses are suppressed during active tasks or locomotion (Otazu et al., 2011; Schneider et al. 2014).

The marmoset is becoming an increasingly popular animal model for systems neuroscience. Their small size, relatively fast generation time, captive breeding success, and sequenced genome (Worley et al., 2014) have made genetic manipulations tractable (e.g., Sasaki et al., 2009). At the same time, it is becoming increasingly possible to adapt the natural behavior of marmosets to a laboratory setting (for example, Miller and Wang, 2006; Roy et al., 2011; Takahashi et al., 2013), and to train marmosets in simple operant behaviors in the auditory (Osmanski et al., 2011) and visual domains (Mitchell et al., 2014). The evolutionary proximity marmosets to other primate species including humans, combined with their genetic and behavioral tractability, thus offers a potent model system to advance our understanding of brain structure and function. Being animals that exhibit a rich vocal behavior and with a well-studied auditory system, the above advantages make marmosets especially attractive to study the brain’s vocal communication machinery. In this study, we have demonstrated an addition to the set of tools that could be used to localize brain regions of higher auditory cortical function and to investigate the processing of communication sounds in this exciting model system.

## Materials and Methods

### Animal preparation

All experimental protocols were approved by the Institutional Animal Care and Use Committees (IACUC) of The Rockefeller University and Weill Cornell Medical College, and met the guidelines of the National Institutes of Health for the care and use of laboratory animals. Six male marmosets (*Callithrix jacchus*), weighing 325 – 400g were imaged while anesthetized. Experimental imaging sessions lasted 90 – 120 min., and each subject was restricted to one imaging session per week. Sedation was induced with intramuscular injection of a combination of ketamine (10mg/kg), dexmedetomidine (0.01mg/kg), and glycopyrrolate (0.005-0.01 mg/kg). Anesthesia was maintained for the duration of MRI imaging using isoflurane (0.5% - 1.5%) in combination with a 50/50 mixture of nitrous oxide (N_2_O) and medical grade oxygen (O_2_), delivered at a rate of 1L/min through a custom-built face mask. Heart rate, core body temperature and respiratory rate were monitored throughout the duration of the imaging procedure using MR-compatible sensors (SA Systems Inc.). Following anesthetic induction, a sterile ophthalmic lubricant was applied to cover both eyes. The anesthetized subject was then placed in the sphinx position within a custom-built anatomic positioner (Figure 1) equipped with built-in circulating warm-water heat support. Earphones for auditory stimulation were placed, and a surface ring coil for functional imaging was positioned over the subjects head. The subject was secured in place using acoustic isolation foam blocks. A custom foam-lined 3D-printed helmet was used to secure the subject’s head and the surface ring coil to the anatomic positioner. About 5 minutes prior to functional imaging, the concentration of isoflurane administered was reduced (0.25 – 0.75%) while concurrently increasing the ratio of N_2_O:O_2_ to 70/30. Isoflurane was kept at low levels during functional imaging in order to maintain responsivity in the auditory cortex. This level of anesthesia is lower than dosages typically used in invasive neurophysiological experiments. At the end of the imaging session, lactated Ringers solution was injected subcutaneously to provide fluid support. The subject recovered from anesthesia in a temperature- and humidity-controlled incubation chamber while under continuous monitoring.

### Stimuli

Stimuli for vocalization experiments consisted of marmoset vocalizations (V), heterospecific vocalizations (H), and phase-scrambled marmoset vocalizations (N, for ‘noisy’). A corpus of 16 non-familiar marmoset vocalizations was constructed, eight from recordings in a colony of marmosets at Johns Hopkins University (Agamaite and Wang, 1997), and the remainder downloaded from various online sources. A corpus of 16 heterospecific vocalizations included calls of other New World primates (tamarins and squirrel monkeys), birds, macaques and other animals, which were downloaded from various online sources. All vocalization stimuli were digitized at 44.1 KHz. Phase-scrambled vocalizations were made by first extracting the power spectrum of marmoset vocalizations in each band of a logarithmic filter bank consisting of 6 equally spaced bands, scrambling their phases, and recombining the scrambled-phase signals from all bands. The average power spectra for the stimulus categories are plotted in Figure 2C. For tone experiments, we generated random-chord stimuli in MATLAB as follows: two frequency bands of two-octave widths, corresponding to low frequencies (center frequency = 1200 Hz) and high frequencies (center frequency = 7000 Hz) were defined, and 21 logarithmically spaced 50 ms-long tone pips were generated in each band. The total stimulus duration was divided into 50 ms bins, and each bin was populated by the sum of randomly drawn tone pips. Tone pips were drawn from the low- or high-frequency band for the low-tone and high-tone stimuli. Tone pip density was maintained constant at ∼5 tone pips/second. We constructed 10 2.25 s-long stimuli in each frequency band, and all 10 were combined in random order to produce a 22.5 s stimulus for each ON block. All acoustic stimuli were normalized to equal broadband power, amplified with a STAX amplifier and presented through MRI-compatible earphones (STAX). Sound level for stimulus presentation was optimized during pilot experiments to maximize the magnitude of the average BOLD response, and the average sound pressure level (measured at a 1-cm distance from the earphone) was ∼80 dB SPL. To reduce ambient scanner noise, the scanner’s helium compressor was switched off during the auditory fMRI runs. An increase in helium gas pressure was observed, but for our short-duration functional scans, this increase was within safe operating limits. We caution that this step may not be appropriate for longer imaging runs.

### Imaging

fMRI was performed on a 7.0 Tesla 70/30 Bruker Biospec small-animal MRI system equipped with a 12 cm diameter, 450 mT/m amplitude, 4500 T/m/s slew rate, actively-shielded gradient subsystem with integrated shim capability. A 3 cm-wide ring surface coil was used for reception of the MR signal and a linear coil with 7 cm diameter was used for the excitation of the sample. In a typical session, after initial localizer and waterline scans, we acquired anatomical images covering the whole brain with a slice thickness of 1.2 mm (18 slices) using a FLASH pulse sequence. Four averages with a flip angle of 40 degrees, TE = 5.5 ms, TR = 355 ms, field of view = 4 × 4 cm, and matrix size = 320 × 320, resulting in a spatial resolution of 0.125 mm × 0.125 mm were acquired. Based on these images, we identified the location of the lateral sulcus (LS), and positioned a slice packet consisting of six 1.2 mm-thick slices parallel to the LS, abutting the LS in the first slice, and extending over the temporal lobe. A second anatomical scan was acquired with this slice prescription using the same parameters as above for future registration with functional images. After completion of anatomical imaging, we commenced functional (EPI) imaging. Six gradient echo EPI image slices of 1.2 mm thickness paralleling the LS as above were acquired with interleaved slice order, TE = 16 ms, flip angle = 80 degrees, navigator echo, field of view = 4 × 4 cm, matrix size = 64 × 64, resulting in an in-plane spatial resolution of 0.625 mm × 0.625 mm. The actual time required for slice acquisition was < 250 ms, but we triggered slice acquisition only every 2.25 s because of our sparse imaging protocol, resulting in an effective TR of 2.25 seconds (Figure 2). We obtained 600 volumes over the course of 22.5 minutes for vocalization experiments, and 400 volumes over 15 minutes for tone experiments. Each run corresponded to 10 repetitions of each stimulus block, with each block lasting 10 TRs, or 22.5 seconds. Each run was preceded by the acquisition of three volumes to overcome the T1 saturation artifact, and these volumes were dropped from analysis.

### Data pre-processing

Data were analyzed using custom scripts in AFNI (Cox, 1996) and MATLAB. We first processed the anatomical volumes by digital skull-stripping and intensity normalization in AFNI. We constructed a mask volume based on the anatomical volume, which used the same slice prescription as the functional volume. We then calculated a mean functional volume over all time points of the functional imaging run, and registered each acquired volume using affine transformations, to the mean functional volume. We saved the affine transformation matrix and motion parameters from this initial registration, but did not resample the functional volumes at this stage. The mean functional maps were then skull-stripped manually and intensity-normalized. We performed a nonlinear alignment of the skull-stripped mean functional to the anatomical volume to remove the effect of EPI distortions. This calculated warp field, together with the affine transformations obtained at the first registration step, was applied to the individual functional volumes to bring them into alignment with the anatomical volume. We de-spiked the resultant volume, and used a list of outliers (points exceeding a threshold of six standard deviations from the mean baseline intensity), to build a list of time frames excluded as likely motion artifacts. The dataset was used for further analysis only if >95% of the time points survived exclusion. The de-spiked volume was high-pass filtered with a cutoff frequency of 0.01 Hz, spatially smoothed using a 2 mm Gaussian kernel, quadratically de-trended, and normalized to a voxel-wise baseline value of 100.

### Data analysis

A general linear model (GLM) was fit (Friston et al., 1995) to these pre-processed data. In analyzing the BOLD responses to both tones and vocalizations, we observed a diversity of response shapes across the subjects. In some cases (Subjects C and P), the BOLD response could be reasonably well-modeled by a standard univariate hemodynamic response function. However, in other subjects (Subjects Y and Q), the BOLD response exhibited two distinct peaks, such that a univariate response model was inadequate to model the observed responses. We therefore adopted a model-free approach for analyzing BOLD activity in our experiments (similar to Gonzalez-Castillo et al., 2012), by using 7-point tent basis functions to fit the shape of the BOLD response over 14 data points in order to accurately account for the observed heterogeneity in the shapes of the hemodynamic response functions. We further observed that the second peak of the BOLD response occurred after stimulus offset, and compared to the first response peak, its magnitude was less modulated stimulus type. These features of the offset response made it difficult to interpret and directly link to underlying neural activity. Therefore, we only used the portion of the response that occurred during the presentation of the stimulus for all the analyses that we have presented here. The design matrix included polynomials up to order two and the six independent affine motion parameters as nuisance regressors. Results were visualized in AFNI, 3DSlicer, and using custom MATLAB scripts. In all cases, we defined regions of interest (ROIs) consisting of voxels where the average beta value across all stimulus types was significantly different from baseline (FDR-corrected q-value = 0.05). From these voxels, we extracted the average time-course of the BOLD response for display. We also obtained beta value maps and maps of the t-statistic from each experimental run.

### Group analysis

To combine data across runs, we first defined a matrix that outlined a 4-voxel (2.5 mm) deep region of the edges of the volume (see description in Results section), stretching 15 mm along the LS from the temporal pole on all six slices and in each hemisphere. In defining this matrix, we explicitly excluded one voxel at the outermost edges of the volume to ensure minimal effects of movement artifacts and partial-volume effects. This resulted in a 24 × 12 matrix, each element of which was the average of the beta values from 4 voxels, and which could be indexed in terms of its anatomical distance from the temporal pole and the LS. Because the brain sizes of the imaged animals were similar, applying this matrix to all subjects allowed us to combine activity across subjects without introducing further registration-induced distortion, partial volume or resampling errors to the functional volumes. We extracted this matrix of betas from individual runs, and fit a second-level GLM to the data, including predictors for each hemisphere and subject. The resulting matrix was then projected back onto an example anatomical scan to map regions of interest in the brain.

## Acknowledgements

We thank Dr. Winrich Freiwald (The Rockefeller University) for support, advice and discussions. We are indebted to Dr. David Troilo (SUNY College of Optometry) for his generous gift of the experimental animals and advice, and to Ann Nour (SUNY College of Optometry) for help with pilot experiments. We are grateful to Dr. Xiaoqin Wang (Johns Hopkins University) for providing some of the marmoset vocalizations used in the study. We thank Skye Rasmussen, DVM and Leslie Diaz, DVM for providing veterinary support during MRI imaging sessions. We thank Drs. Winrich Freiwald and Xiaoqin Wang for critical comments on the manuscript. This material is based upon work supported by the Center for Brains, Minds and Machines (CBMM), funded by NSF STC award CCF-1231216, and funds from The Rockefeller University. Srivatsun Sadagopan was supported by a Leon Levy postdoctoral fellowship. Henning Voss was funded by NSF award #0956306 and the Nancy M. and Samuel C. Fleming Research Scholar Award in Intercampus Collaborations.

## Author contributions

SS and HUV designed the experiments. HUV designed the MRI sequences and the gaseous anesthesia delivery system, provided advice on stimulus delivery and data analysis and assisted with imaging. SS and NZT collected and analyzed data. SS wrote the main manuscript text, and all authors reviewed the manuscript.

## References

Agamaite JA & Wang X (1997) Quantitative classification of the vocal repertoire of the common marmoset, Callithrix jacchus. Master’s thesis, The Johns Hopkins University, Baltimore MD.

Aitkin LM, Merzenich MM, Irvine DR, Clarey JC & Nelson JE (1986). Frequency representation in the auditory cortex of the common marmoset (Callithrix jacchus jacchus). J Comp Neurol 252: 175–185.

Aitkin L & Park V (1993). Audition and the auditory pathway of a vocal New World primate, the common marmoset. Prog Neurobiol 41: 345–367.

Allen EA, Erhardt EB & Calhoun VD (2012). Data visualization in the neurosciences: overcoming the curse of dimensionality. Neuron 74: 603–608.

Andics A, Gacsi M, Farago T, Kis A & Miklosi A (2014). Voice-sensitive regions in the dog and human brain are revealed by comparative fMRI. Curr Biol 24: 574–8.

Belin P, Zatorre RJ, Lafaille P, Ahad P & Pike B (2000). Voice-selective areas in human auditory cortex. Nature 403: 309–312.

Bendor D & Wang X (2008). Neural response properties of primary, rostral, and rostrotemporal core fields in the auditory cortex of marmoset monkeys. J Neurophysiol 100: 888–906.

Chaplin TA, Yu HH, Soares JG, Gattass R & Rosa MP (2013). A conserved pattern of differential expansion of cortical areas in simian primates.

Cox RW (1996). AFNI: Software for the analysis and visualization of functional magnetic resonance neuroimages. Computers and Biomedical Research 29: 162–73.

DiMattina C & Wang X (2006). Virtual vocalization stimuli for investigating neural representations of species-specific vocalizations. J Neurophysiol 95: 1244–62.

Dunbar RIM (1992). Neocortex size as a constraint on group size in primates. J Hum Evol 20: 469–493.

Epple G (1968). Comparative studies on vocalization in marmoset monkeys (Hapalidae). Folia Primatol 8: 1–40.

Friston KJ, Holmes AP, Worley KJ, Poline JP, Frith CD & Frackowiak RSJ (1995). Statistical parametric maps in functional imaging: a general linear approach. Human Brain Mapping 2: 189–210.

Fukushima M, Saunders RC, Leopold DA, Mishkin M & Averbeck BB (2014). Differential coding of conspecific vocalizations in the ventral auditory cortical stream. J Neurosci 34: 4665–76.

Gaese BH & Ostwald J (2001). Anesthesia changes frequency tuning of neurons in the rat primary auditory cortex. J Neurophysiol 86: 1062–6.

Gonzalez-Castillo J, Saad ZS, Handwerker DA, Inati SJ, Brenowitz N & Bandettini PA (2012). Wholebrain, time-locked activation with simple tasks revealed using massive averaging and model-free analysis. Proc. Natl. Acad. Sci. USA 109: 5487–5492.

Goodman M, Porter CA, Czelusniak J, Page SL, Schneider H, Shoshani J, Gunnell G & Groves CP (1998). Toward a phylogenetic classification of primates based on DNA evidence complemented by fossil evidence. Mol Phylogenet Evol 9: 585–98.

Grimsley JM, Shanbhag SJ, Palmer AR & Wallace MN (2012). Processing of communication calls in Guinea pig auditory cortex. PLoS One 7: e51646.

Hall AJ, Brown TA, Grahn JA, Gati JS, Nixon PL, Hughes SM, Menon RS & Lomber SG (2014). There’s more than one way to scan a cat: imaging cat auditory cortex with high-field fMRI using continuous or sparse sampling. J Neurosci Methods 224: 96–106.

Holmstrom LA, Eeuwes LB, Roberts PD & Portfors CV (2010). Efficient encoding of vocalizations in the auditory midbrain. J Neurosci 30: 802–19.

Issa EB, Papanastassiou AM & DiCarlo JJ (2013). Large-scale, high-resolution neurophysiological maps underlying FMRI of macaque temporal lobe. J Neurosci 33: 15207–19.

Kadia SC & Wang X (2003). Spectral integration in A1 of awake primates: neurons with single- and multipeaked tuning characteristics. J Neurophysiol 89: 1603–22.

Kajikawa Y, de la Mothe L, Blumell S & Hackett TA (2005). A comparison of neuron response properties in areas A1 and CM of the marmoset monkey auditory cortex: tones and broadband noise. J Neurophysiol 93: 22–34.

Kajikawa Y, de la Mothe LA, Blumell S, Sterbing-D’Angelo SJ, D’Angelo W, Camalier CR & Hackett TA (2008). Coding of FM sweep trains and twitter calls in area CM of marmoset auditory cortex. Hear Res 239: 107–25.

Kriegstein KV & Giraud AL (2004). Distinct functional substrates along the right superior temporal sulcus for the processing of voices. Neuroimage 22: 948–55.

Ku SP, Tolias AS, Logothetis NK & Goense J (2011). fMRI of the face-processing network in the ventral temporal lobe of awake and anesthetized macaques. Neuron 70: 352–62.

Liu JV, Hirano Y, Nascimento GC, Stefanovic B, Leopold DA & Silva AC (2013). fMRI in the awake marmoset: somatosensory-evoked responses, functional connectivity, and comparison with propofol anesthesia. Neuroimage 78:186–95.

Miller CT & Wang X (2006). Sensory-motor interactions modulate a primate vocal behavior: antiphonal calling in common marmosets. J Comp Physiol A Neuroethol Sens Neural Behav Physiol 192: 27–38.

Mitchell JF, Reynolds JH & Miller CT (2014). Active vision in marmosets: a model system for visual neuroscience. J Neurosci 34: 1183–94.

de la Mothe LA, Blumell S, Kajikawa Y & Hackett TA (2006). Cortical connections of the auditory cortex in marmoset monkeys: core and medial belt regions. J Comp Neurol 496: 27–71.

de la Mothe LA, Blumell S, Kajikawa Y & Hackett TA (2012). Cortical connections of auditory cortex in marmoset monkeys: lateral belt and parabelt regions. Anat Rec (Hoboken) 295: 822–36.

Nagarajan SS, Cheung SW, Bedenbaugh P, Beitel RE, Schreiner CE & Merzenich MM (2002). Representation of spectral and temporal envelope of twitter vocalizations in common marmoset primary auditory cortex. J Neurophysiol 87: 1723–37.

Osmanski MS & Wang X (2011). Measurement of absolute auditory thresholds in the common marmoset (Callithrix jacchus). Hear Res 277: 127–33.

Otazu GH, Tai LH, Yang Y & Zador AM (2009). Engaging in an auditory task suppresses responses in auditory cortex. Nat Neurosci 12: 646–54.

Papoti D, Yen CC, Mackel JB, Merkle H & Silva AC (2013). An embedded four-channel receive-only RF coil array for fMRI experiments of the somatosensory pathway in conscious awake marmosets. NMR Biomed 26: 1395–402.

Perrodin C, Kayser C, Logothetis NK & Petkov CI (2011). Voice cells in the primate temporal lobe. Curr Biol 21: 1408–15.

Petkov CI, Kayser C, Augath M & Logothetis NK (2006). Functional imaging reveals numerous fields in the monkey auditory cortex. PLoS Biol 4: e215.

Petkov CI, Kayser C, Steudel T, Whittingstall K, Augath M & Logothetis NK (2008). A voice region in the monkey brain. Nat Neurosci 11: 367–74.

Petkov CI, Logothetis NK & Obleser J (2009). Where are the human speech and voice regions, and do other animals have anything like them? Neuroscientist 15: 419–29.

Petkov CI, Kayser C, Augath M & Logothetis NK (2009). Optimizing the imaging of the monkey auditory cortex: sparse vs. continuous fMRI. Magn Reson Imaging 27: 1065–73.

Philibert B, Beitel RE, Nagarajan SS, Bonham BH, Schreiner CE & Cheung SW (2005). Functional organization and hemispheric comparison of primary auditory cortex in the common marmoset (Callithrix jacchus). J Comp Neurol 487: 391–406.

Pistorio AL, Vintch B & Wang X (2006) Acoustic analysis of vocal development in a New World primate, the common marmoset (Callithrix jacchus). J Acoust Soc Am 120: 1655–70.

Pollack GD (2013). The dominant role of inhibition in creating response selectivities for communication calls in the brainstem auditory system. Hear Res 305: 86–101.

Portfors CV, Roberts PD & Jonson K (2009). Over-representation of species-specific vocalizations in the awake mouse inferior colliculus. Neuroscience 162: 486–500.

Rauschecker JP & Tian B (2000). Mechanisms and streams for processing of “what” and “where” in auditory cortex. Proc Natl Acad Sci USA 97: 11800–6.

Recanzone GH, Guard DC & Phan ML (2000). Frequency and intensity response properties of single neurons in the auditory cortex of the behaving macaque monkey. J Neurophysiol 83: 2315–31.

Remedios R, Logothetis NK & Kayser C (2009). An auditory region in the primate insular cortex responding preferentially to vocal communication sounds. J Neurosci 29: 1034–45.

Rojas MJ, Navas JA, Greene SA & Rector DM (2008). Discriminiation of auditory stimuli during isoflurane anesthesia. Comp Med 58: 454–7.

Romanski LM & Averbeck BB (2009). The primate cortical auditory system and neural representation of conspecific vocalizations. Annu Rev Neurosci 32: 315–46.

Roy S, Miller CT, Gottsch D & Wang X (2011). Vocal control by the common marmoset in the presence of interfering noise. J Exp Biol 214: 3619–29.

Sadagopan S & Wang X (2009). Nonlinear spectrotemporal interactions underlying selectivity for complex sounds in auditory cortex. J Neurosci 29: 11192–202.

Sasaki E et al. (2009). Generation of transgenic non-human primates with germline transmission. Nature 459: 523–7.

Schneider DM, Nelson A & Mooney R (2014). A synaptic and circuit basis for corollary discharge in the auditory cortex. Nature 513: 189–94.

Scott BH, Malone BJ & Semple MN (2007). Effect of behavioral context on representation of a spatial cue in core auditory cortex of awake macaques. J Neurosci 27: 6489–99.

Scott SK, Blank CC, Rosen S & Wise RJS (2000). Identification of a pathway for intelligible speech in the left temporal lobe. Brain 123: 2400–2406.

Snowdon CT (2001). Social processes in communcation and cognition in callitrichid monkeys: a review. Anim Cogn 4: 247–57.

Steiper ME & Young NM (2006). Primate molecular divergence dates. Mol Phylogenet Evol 41: 384–394.

Stevenson MF & Poole TB (1976). An ethogram of the common marmoset (Callithrix jacchus jacchus): general behavioral response. Anim Behav 24: 428–451.

Suta D, Kvasnak E, Popelar J & Syka J (2003). Representation of species-specific vocalizations in the inferior colliculus of the guinea pig. J Neurophysiol 90: 3794–808.

Taglialatela JP, Russell JL, Schaeffer JA & Hopkins WD (2009). Visualizing vocal perception in the chimpanzee brain. Cereb Cortex 19: 1151–7.

Takahashi DY, Narayanan DZ & Ghazanfar AA (2013). Coupled oscillator dynamics of vocal turn-taking in monkeys. Curr Biol 23: 2162–8.

Ter-Mikaelian M, Sanes DH & Semple MN (2007). Transformation of temporal properties between auditory midbrain and cortex in the awake Mongolian gerbil. J Neurosci 27: 6091–102.

Worley KC et al. (2014). The common marmoset genome provides insight into primate biology and evolution. Nat Genetics 46: 850–857.

Tian B, Reser D, Durham A, Kustov A & Rauschecker JP (2001). Functional specialization in rhesus monkey auditory cortex. Science 292: 290–3.

Tsao DY, Freiwald WA, Knutsen TA, Mandeville JB & Tootell RB (2003). Faces and objects in macaque cerebral cortex. Nat Neurosci 6: 989–95.

Tsao DY, Freiwald WA, Tootell RB & Livingstone MS (2006). A cortical region consisting entirely of face-selective cells. Science 311: 670–4.

Wang X, Merzenich MM, Beitel R & Schreiner CE (1995). Representation of a species-specific vocalization in the primary auditory cortex of the common marmoset: temporal and spectral characteristics. J Neurophysiol 74: 2685–706.

Wang X (2000). On cortical coding of communication sounds in primates. Proc Natl Acad Sci USA 97: 11843–9.

Wang X & Kadia SC (2001). Differential representation of species-specific primate vocalizations in the auditory cortices of marmoset and cat. J Neurophysiol 86: 2616–20.

Wang X, Lu T, Snider RK & Liang L (2005). Sustained firing in auditory cortex evoked by preferred stimuli. Nature 435: 341–6.

Zhou Y & Wang X (2014). Spatially extended forward suppression in primate auditory cortex. Eur J Neurosci 39: 919–33.

